# Yap is a key regulator of tracheoesophageal separation during foregut development

**DOI:** 10.64898/2026.11.01.763315

**Authors:** Eugene Kim, Jianwen Que

**Affiliations:** Lung Biology Laboratory, Division of Pulmonary, Allergy, and Critical Care Medicine, Department of Medicine, Vagelos College of Physicians and Surgeons, Columbia University Irving Medical Center, New York, NY, USA; Columbia Center for Human Development and Division of Digestive and Liver Diseases, Department of Medicine, Vagelos College of Physicians and Surgeons, Columbia University Irving Medical Center, New York, NY, USA

## Abstract

Separation of the anterior foregut into the esophagus and trachea involves dynamic cellular movement that requires the coordinated activity of multiple signaling pathways and transcription factors. Disruption of this process can result in esophageal atresia (EA) with or without tracheoesophageal fistula (TEF), manifesting in five major anatomical types (Types A-E). Here, we examine the role of the Hippo pathway effector Yes- associated protein (Yap) in foregut separation. Conditional deletion of *Yap* in the dorsal and medial portions of the anterior foregut endoderm leads to the formation of Type A EA/TEF, isolated EA. This separation defect only occurs in *Sox2-CreER;Yap^loxp/loxp^* but not in *Shh-Cre;Yap^loxp/loxp^* mutants. Our findings further reveal that *Yap* mutants exhibit a reduction in Sox2^+^ esophageal progenitor cells accompanied by the absence of the middle esophageal segment. Combined deletion of *Yap* and *Taz* (transcriptional coactivator with PDZ-binding motif) also results in isolated EA accompanied by severe craniofacial abnormalities. Therefore, loss of *Yap* causes a distinct Type A EA/TEF, which differs from existing mouse models of EA/TEF that predominantly exhibit the Type C phenotype. Our findings also suggest that epithelial cells at the dorsal-ventral boundary play a critical role in tracheoesophageal separation.

## Introduction

The esophagus and trachea arise from a single foregut tube during embryonic development. The dorsal side of the anterior foregut gives rise to the esophagus, and the ventral side of the foregut forms the trachea (Kluth and Fiegel, 2003; O’Rahilly and Müller, 1984). Abnormal separation of the esophagus and trachea results in esophageal atresia with or without tracheoesophageal fistula (EA/TEF) (Haight, 1944), a congenital anomaly occurring in approximately 1 in 3,500 live births (Malmström et al., 2008; Spitz, 2007; Torfs et al., 1995). Based on the Gross classification, EA/TEF is categorized into five anatomical types (Types A-E), each characterized by distinct patterns of esophageal discontinuity and tracheoesophageal communication (Fig. 1A). Although multiple genetic contributors have been identified, the etiology of EA/TEF is poorly understood.

**Fig. 1.**
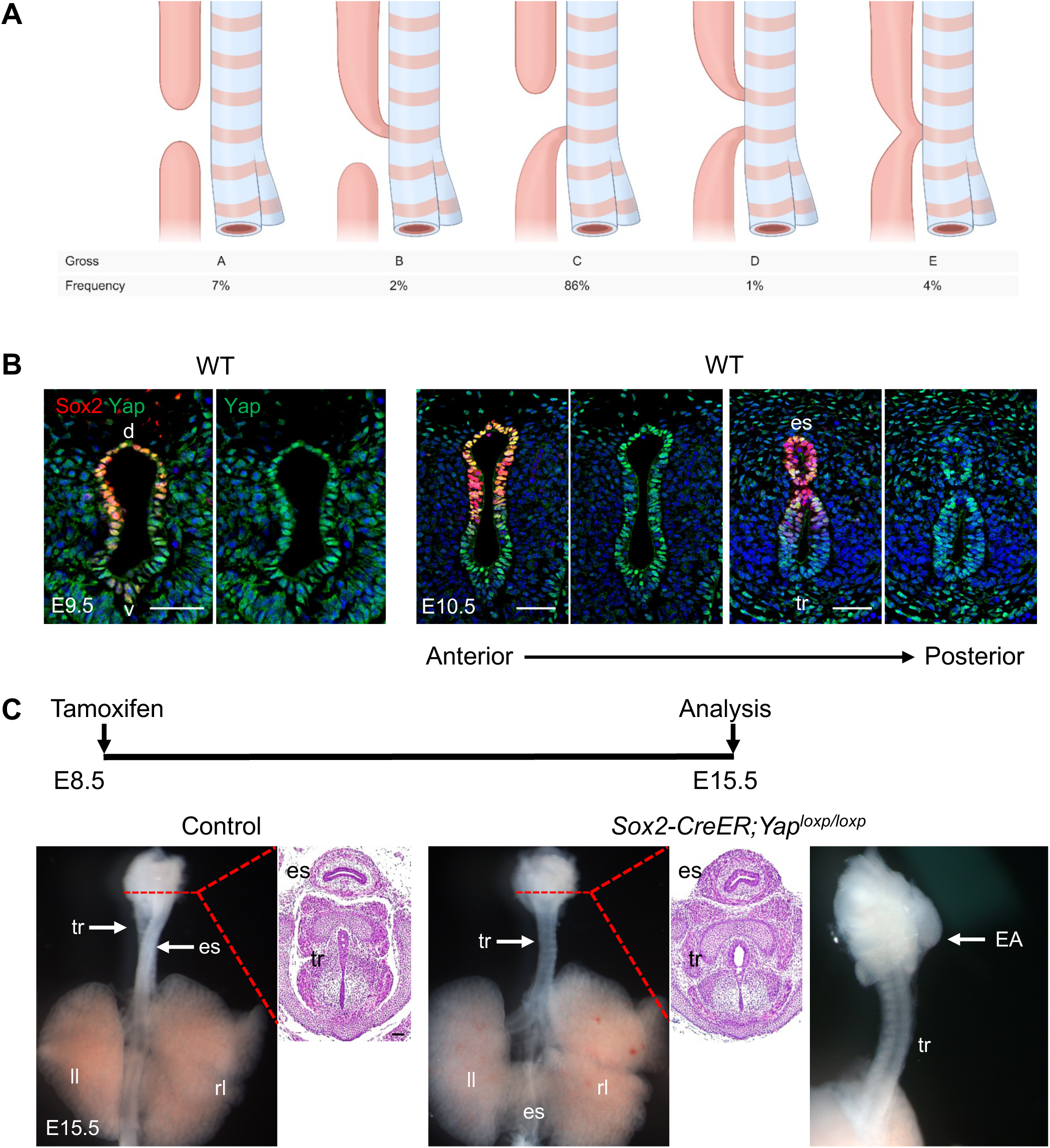
*Yap* deletion leads to the formation of isolated esophageal atresia (Type A EA/TEF) in *Sox2-CreER;Yap^loxp/loxp^*mutant embryos. (A) The five anatomical types of EA/TEF according to the Gross classification and their relative frequencies. Esophagus, pink; trachea, blue. Redrawn and adapted from Krishnan (2019) under a CC BY 4.0 license. (B) Immunofluorescence images of transverse sections from the foreguts of wild-type embryos at E9.5 and E10.5. The dorsal epithelium is marked by Sox2 expression. Note that nuclear Yap expression is abundant throughout the foregut epithelial cells before and during tracheoesophageal separation. (C) Isolated esophageal atresia in *Sox2-CreER;Yap^loxp/loxp^* mutants at E15.5. Note the intact respiratory tract, with esophageal atresia at the proximal end and a separate distal esophageal segment that remains connected to the stomach (data not shown) (n=5). H&E staining of transverse sections from control and *Yap* mutant embryos indicates the separation of the trachea and esophagus in the anterior region. d, dorsal; EA, esophageal atresia; es, esophagus; ll, left lung; rl, right lung; tr, trachea; v, ventral; WT, wild type. Scale bars: 50 μm.

In mice, separation of the esophagus and trachea begins around embryonic day (E) 9.5 and is completed by E11.5 (Cardoso and Lü, 2006; Warburton et al., 2000). Tracheoesophageal separation involves dynamic cellular rearrangements, including coordinated movements of epithelial and mesenchymal cells, as the respiratory and digestive tracts develop (Kim et al., 2019; Yan et al., 2025). Notably, a small number of respiratory progenitor cells expressing Nkx2.1 intercalate into the ventral esophageal epithelium, while esophageal epithelial cells may reciprocally contribute to the dorsal tracheal epithelium, suggesting bidirectional epithelial cell incorporation during foregut separation (Kim et al., 2019). Moreover, previous studies have shown that transcription factors such as Sox2, Nkx2.1, and Isl1 are expressed in the anterior foregut endoderm, and the loss of these genes results in defects in foregut separation (Que et al., 2007; Wood and Episkopou, 1999; Minoo et al., 1999; Jacobs et al., 2012; Kim et al., 2019). Notably, these transcription factors exhibit distinct dorsoventral expression patterns within the developing foregut. For example, Sox2 is highly expressed in the dorsal foregut endoderm, and *Sox2* hypomorphic mouse mutants develop EA/TEF, highlighting the importance of precise transcriptional regulation in tracheoesophageal development (Que et al., 2007).

Similarly, components and activity of several key signaling pathways, including Wnt, Sonic hedgehog (Shh), and bone morphogenetic protein (BMP), exhibit distinct dorsoventral patterns of expression and activity during foregut development. Disruption of these signaling pathways also impairs normal tracheoesophageal separation, contributing to the development of EA/TEF (Goss et al., 2009; Harris-Johnson et al., 2009; Litingtung et al., 1998; Pepicelli et al., 1998; Geneviève et al., 2007; Motoyama et al., 1998; Mahlapuu et al., 2001; Domyan et al., 2011; Li et al., 2007; Que et al., 2006). However, whether the Hippo signaling pathway, its effector Yap in particular, contributes to tracheoesophageal separation remains unclear.

The Hippo signaling pathway is a key regulator of organ size and tissue homeostasis, controlling cell proliferation and apoptosis (Pan, 2010). The core components of the Hippo pathway include a kinase cascade and transcriptional coactivators. In mammals, activation of the Hippo pathway leads to phosphorylation of the transcriptional coactivators Yap and Taz (Zhao et al., 2007; Lei et al., 2008). Phosphorylated Yap and Taz bind to 14-3-3 proteins, resulting in their retention in the cytoplasm and subsequent ubiquitin dependent degradation following further phosphorylation by casein kinase 1 (CK1) (Zhao et al., 2007; Zhao et al., 2010). Conversely, when the Hippo pathway is inactivated, unphosphorylated Yap and Taz translocate into the nucleus, where they interact with TEAD family transcription factors (TEAD1-4) to activate the expression of genes that promote cell growth and proliferation (Zhao et al., 2008).

Homozygous *Yap*-null embryos fail to develop beyond approximately E8.5, before tracheoesophageal separation, which precludes analysis of the role of Yap in this process (Morin-Kensicki et al., 2006). We previously showed that *Shh-Cre*-mediated *Yap* deletion reduces epithelial progenitor proliferation and stratification in the esophagus without disrupting tracheoesophageal separation (Bailey et al., 2019). Of note is that *Shh-Cre* is active predominantly in the ventral foregut endoderm during tracheoesophageal separation (Rodriguez et al., 2010). Intriguingly, conditional deletion of *Yap* using *Sox2-CreER* in this study results in Type A EA/TEF, a phenotype distinct from the predominantly Type C EA/TEF observed in other available mouse models.

Furthermore, combined deletion of *Yap* and *Taz* does not appear to exacerbate tracheoesophageal separation defects beyond those observed following *Yap* deletion alone. Collectively, these findings suggest that Yap plays a critical role in establishing the boundary between the developing respiratory and esophageal epithelial cells, potentially by regulating epithelial cell intercalation during foregut separation.

## Results and Discussion

### *Yap* deletion in Sox2^+^ foregut endoderm leads to isolated EA in mouse embryos

We first characterized Yap expression during anterior foregut development by performing immunofluorescence staining for Yap in wild-type mouse embryos. Yap exhibited robust expression in both the dorsal and ventral epithelium of the anterior foregut at E9.5 (Fig. 1B). Yap expression in the foregut epithelium was maintained throughout tracheoesophageal separation. At E10.5, as tracheoesophageal separation progresses from caudal to rostral, the rostral foregut is not yet separated. Yap expression was observed in the separated esophagus and trachea as well as in the primary lung buds (Fig. 1B and Fig. S1A). To investigate the role of Yap in tracheoesophageal separation, we utilized *Sox2-CreER* mice to conditionally ablate *Yap* in the medial and dorsal regions of the foregut endoderm. Pregnant females were injected intraperitoneally with tamoxifen (0.1 mg/g body weight) at E8.5 to induce Cre- mediated recombination before foregut separation. In the E10.5 *Sox2-CreER;Yap^loxp/loxp^* mutant foregut, Yap expression was not detected in the dorsal foregut epithelium or the separated esophagus. By contrast, Yap was still detected in the tracheal epithelium (Fig. S1B).

In contrast to *Shh-Cre;Yap^loxp/loxp^* mutants (Bailey et al., 2019), isolated EA (Type A EA/TEF) developed in *Sox2-CreER;Yap^loxp/loxp^* mutants with complete penetrance (n=5). A single tracheal tube was present when examined at E15.5, while the proximal esophagus terminated in a blind pouch (Fig. 1C). The distal esophageal segment remained disconnected from the proximal esophagus (Fig. 1C) but continued to connect to the stomach (data not shown). Hematoxylin and eosin (H&E)-stained sections showed the proximal esophageal segment separated from the trachea (Fig. 1C).

Together, these findings suggest that Yap is required for esophageal development during tracheoesophageal separation. Importantly, the distinct phenotypes resulting from *Shh-Cre-* and *Sox2-CreER*-mediated *Yap* deletion suggest that Yap function in epithelial cells at the dorsoventral boundary of the developing foregut is critical for proper tracheoesophageal separation.

### Tracheal and lung development is largely preserved in *Sox2-CreER;Yap^loxp/loxp^* mutants

To assess tracheal and lung development, we analyzed E18.5 *Yap* mutant and control embryos by histology and immunostaining for cell-type markers. Histological and marker analyses revealed no apparent differences between mutants and controls. Sox2^+^ epithelial cells, p63^+^ basal cells, and Sox9^+^ cartilage were detected in the trachea of *Yap* mutants (Fig. 2A). Moreover, the alveolar type I marker T1α, the ciliated cell marker acetylated α-tubulin (A-tu), the alveolar type II marker Sftpc, and the club cell marker Scgb1a1 were all detected in the mutant lungs (Fig. 2B). These findings suggest that *Yap* deletion prior to tracheoesophageal separation has minimal effects on respiratory system development in *Sox2-CreER;Yap^loxp/loxp^* mutants. Consistent with this observation, *Yap* deletion was largely confined to the dorsal foregut, and Yap expression persisted in the tracheal epithelium when tamoxifen was given at E8.5 (Fig. S1B).

**Fig. 2.**
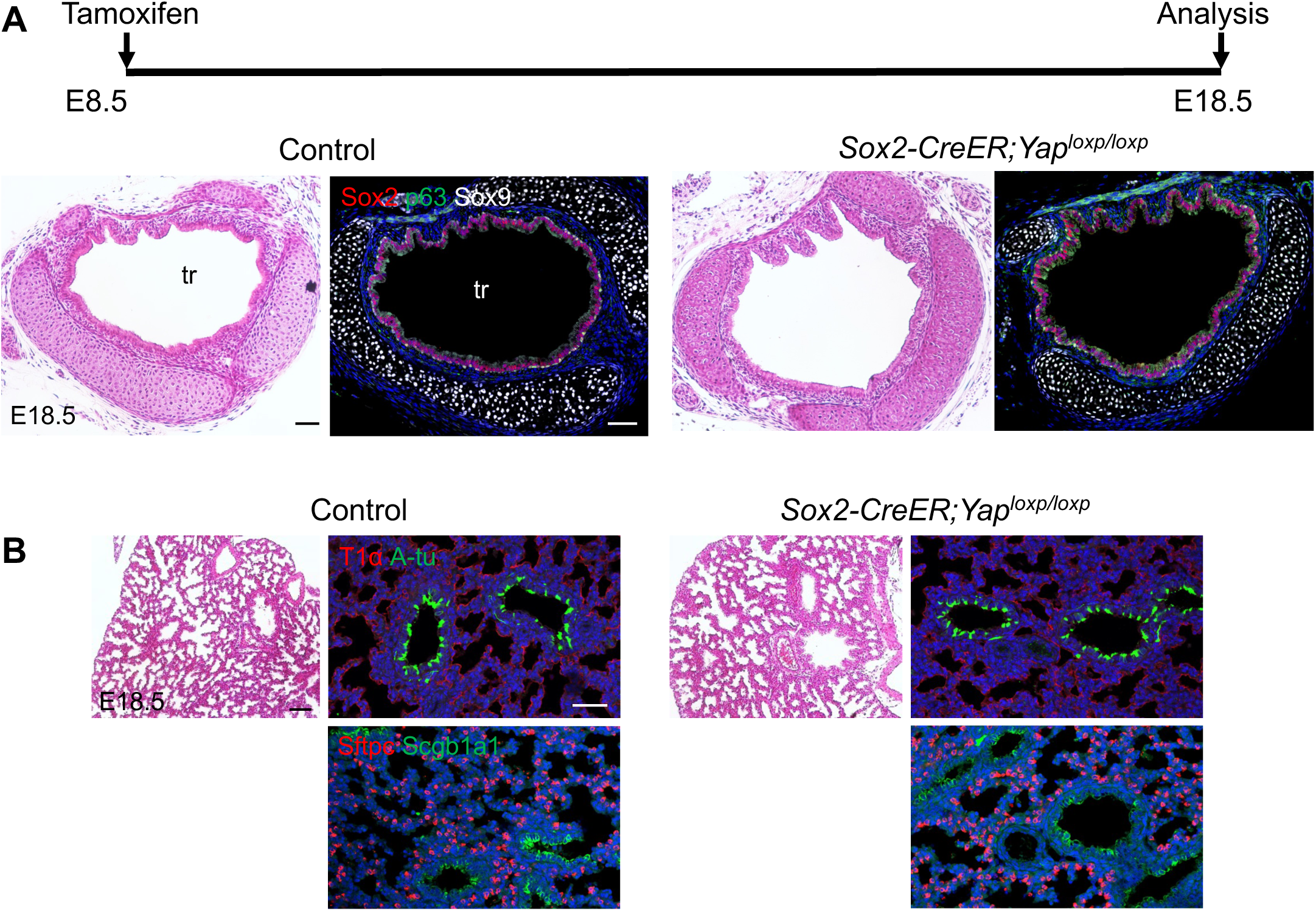
Normal development of the respiratory system in *Sox2-CreER;Yap^loxp/loxp^* mutant embryos. (A) H&E staining and immunofluorescence of transverse sections of the trachea from E18.5 control and *Yap* mutant embryos (n=3). The trachea is lined by a pseudostratified columnar epithelium and contains C-shaped cartilage. Note the presence of Sox2^+^ epithelial cells, p63^+^ basal cells and Sox9^+^ cartilage in the *Yap* mutant trachea. (B) The lungs of E18.5 *Yap* mutants show terminal sac morphology similar to that in control embryos. Airway (Acetylated α-tubulin (A-tu) and Scgb1a1) and alveolar epithelial markers (T1α and Sftpc) are detected in mutant lungs with distributions similar to those in controls. tr, trachea. Scale bars: 50 μm.

### *Yap* deletion reduces esophageal progenitors and leaves ectopic epithelial cells during foregut separation

To investigate the tracheoesophageal separation process in *Yap* mutants, we analyzed mutant embryos at E10.5 and E11.5 by immunostaining with Sox2 and Nkx2.1 to label esophageal and respiratory progenitor cells, respectively. At E10.5, Sox2^+^ cells were located dorsally in the *Yap* mutant foregut, but fewer Sox2^+^Nkx2.1^-^ epithelial cells were observed than in the control at both the tracheal bifurcation and lung-bud levels (Fig. 3A), with sparse Ki67^+^ proliferating cells (Fig. S2A). By contrast, Nkx2.1 expression was retained in both the ventral foregut and the primary lung buds (Fig. 3A). Additionally, p63^+^ progenitors remained present in the dorsal epithelium and separated esophagus of the *Yap* mutant foregut, although fewer were observed per section than in the control (Fig. S2B). These observations suggest that Yap is required for the expansion of Sox2^+^ esophageal progenitors during tracheoesophageal separation.

**Fig. 3.**
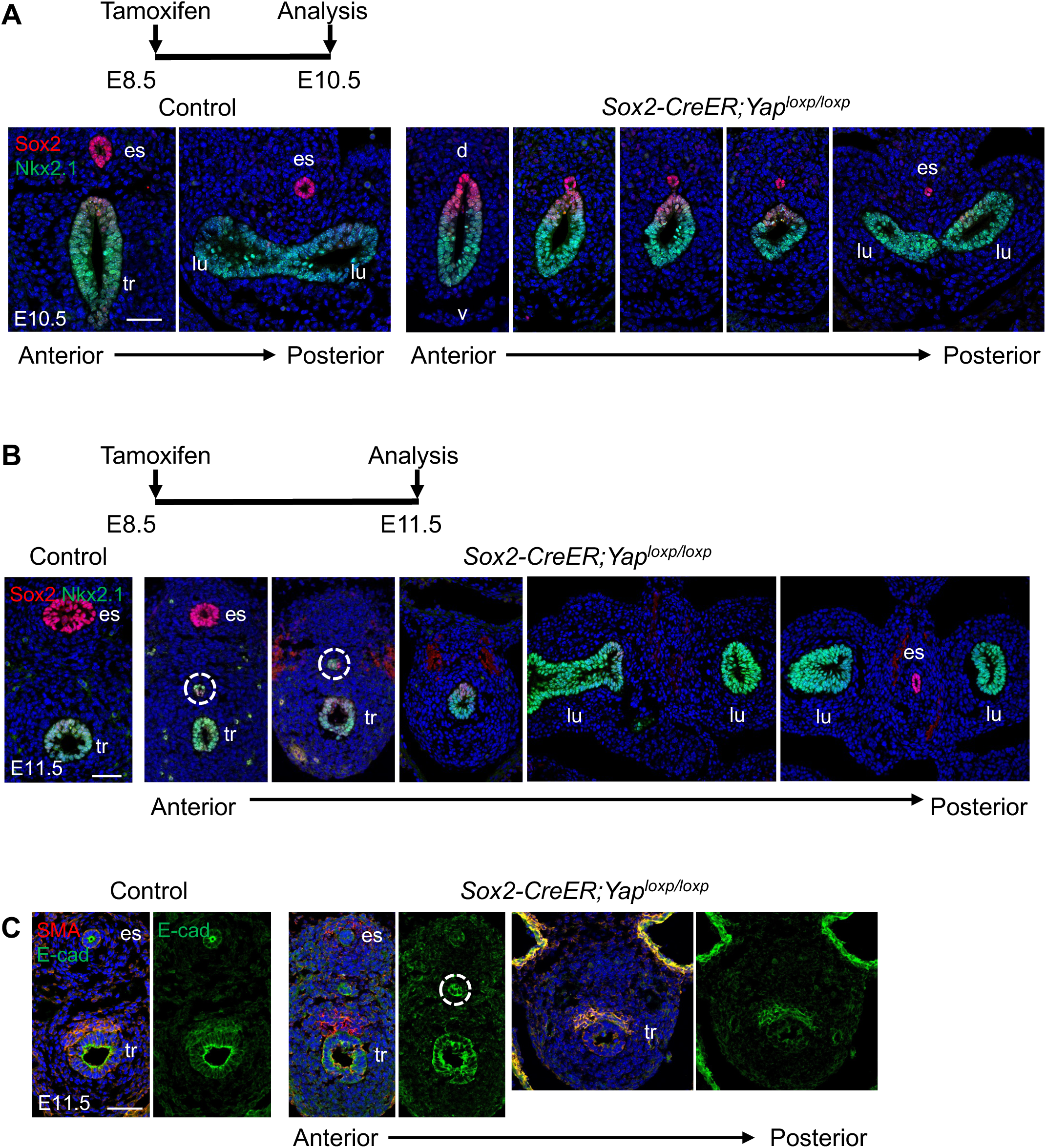
Reduced numbers of Sox2^+^ esophageal epithelial cells and ectopic presence of Sox2^+^Nkx2.1^+^ cells in the foregut of *Sox2-CreER;Yap^loxp/loxp^* mutants. (A-B) Immunofluorescence staining of transverse sections from the *Yap* mutant foregut at E10.5 and E11.5. Sox2 marks esophageal (dorsal) epithelial cells, while Nkx2.1 marks tracheal and lung epithelial cells. (A) The *Yap* mutant foregut at E10.5 contains fewer Sox2^+^ epithelial cells than the control along the anterior-posterior axis. The number of Sox2^+^Nkx2.1^-^ epithelial cells per section is reduced in the mutant at matched axial levels (tracheal bifurcation: control 16.3 ± 3.2, mutant 4.0 ± 1.7; lung bud: control 31.7 ± 10.0, mutant 14.5 ± 3.3; mean ± SD, n = 3-4 sections per level from one embryo per genotype). Control, *Sox2-CreER;Yap^loxp/+^*. (B) Sox2^+^ esophageal epithelial cells are absent in the middle but present in the anterior and posterior regions of the esophagus upon *Yap* deletion. An ectopic cluster of Sox2^+^Nkx2.1^+^ cells (dashed circle) is located between the esophagus and trachea in the anterior foregut. (C) E-cadherin (E-cad) and α-smooth muscle actin (SMA) staining identify epithelial cells and smooth muscle cells, respectively. E-cadherin staining shows that the ectopic cluster forms a circular structure (dashed circle). At E11.5, the esophagus is absent at mid-tracheal and bronchial levels in two mutant embryos (6 bronchial-level sections examined in total) but present in all sections from control embryos. d, dorsal; es, esophagus; lu, lung; tr, trachea; v, ventral. Scale bars: 50 μm.

Serial sections of E11.5 *Yap* mutants from anterior to posterior revealed the absence of the middle esophageal segment, with Sox2^+^ esophageal cells being retained in the anterior and posterior regions (Fig. 3B). E-cadherin immunostaining confirmed the absence of the middle portion of the esophagus (Fig. 3C). Notably, an ectopic cluster of cells was found between the esophagus and trachea in the anterior foregut of the *Yap* mutants. Further posteriorly, the cluster was located in areas adjacent to tracheal epithelial cells in the mutant foregut (Fig. 3B). The ectopic cells co-expressed Sox2 and Nkx2.1 (Fig. 3B), and E-cadherin staining supported their epithelial identity (Fig. 3C).

We also observed an ectopic epithelial tube adjacent to the mutant trachea at E18.5 (Fig. S3A). Histological examination revealed a tube-like structure composed of epithelial and mesenchymal layers (Fig. S3A), and the surrounding mesenchyme lacked cartilage. While Sox2 was expressed in the majority of epithelial cells within the tube, p63 expression was restricted to a smaller subset of these cells (Fig. S3B). Interestingly, a subpopulation in the tube expressed both Nkx2.1 and Foxj1 (Fig. S3C), suggesting that these cells exhibit respiratory epithelial features.

### The proximal and distal esophageal segments are differentially affected in *Yap* mutants

Although the middle portion of the esophagus was absent in *Yap* mutants, the proximal and distal segments remained. To assess the esophageal development of these segments, we performed histological analysis and immunofluorescence staining. The proximal segment at E18.5 was lined with the stratified squamous epithelium, similar to the control esophagus (Fig. 4A). The presence of esophageal progenitor cells was confirmed by immunostaining for Sox2 and p63, which were expressed in the basal layer of the esophageal epithelium (Fig. 4A). The differentiation marker Krt13 was expressed in the suprabasal layers of the proximal esophagus in both controls and *Yap* mutants (Fig. 4B). The columnar cell marker Krt8 was restricted to the luminal layers of the stratified epithelium and was not expressed in the basal or suprabasal layers of either the control or mutant esophagus (Fig. 4B). However, the distal segment of the *Yap* mutant esophagus exhibited disorganized epithelial cell layers with a mixture of squamous and columnar epithelium (Fig. 4C). While Sox2 expression was detected in most epithelial cells, p63^+^ cells were fewer and discontinuous (Fig. 4C), suggesting altered basal cell composition in the residual distal esophagus. Moreover, the distal segment was lined by Krt8^+^ columnar cells, and few Krt13^+^ cells were detected (Fig. 4C), indicating that the distal epithelium retained columnar features and had not acquired normal squamous differentiation by E18.5.

**Fig. 4.**
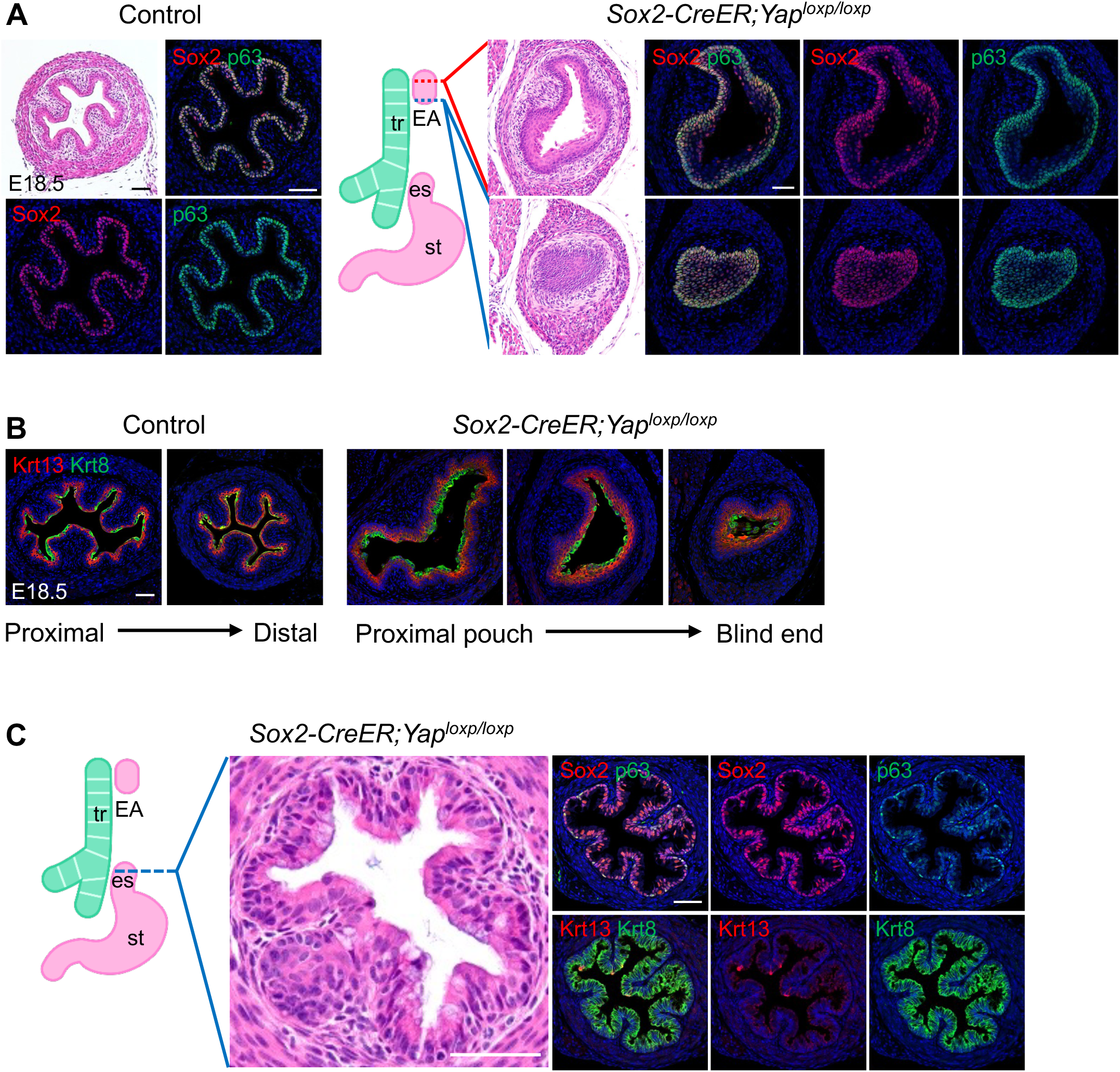
Abnormalities in esophageal development are observed in *Yap* mutants at E18.5. (A) H&E and immunofluorescence staining for Sox2 and p63 in the esophagus of control and *Yap* mutant embryos (n=3). The diagram shows the trachea in green and the proximal esophageal pouch, distal esophageal segment and stomach in pink, with a gap indicating the absent middle esophageal segment. H&E staining of the proximal esophagus in *Yap* mutants reveals a stratified squamous epithelium. Sox2 and p63 are expressed in the basal cells of the proximal esophagus and its blind end in *Yap* mutants. (B) Three sections of the proximal esophageal pouch of a *Yap* mutant, arranged from proximal to distal toward the blind end, show Krt13 in suprabasal cells and Krt8 in luminal cells, similar to the control. (C) H&E staining shows the abnormal epithelial morphology in the distal segment of the *Yap* mutant esophagus. Note Sox2 expression in most epithelial cells, sparse p63 and Krt13 staining and Krt8^+^ columnar epithelial cells in the residual distal segment. EA, esophageal atresia; es, esophagus; st, stomach; tr, trachea. Scale bars: 50 μm.

### Combined deletion of *Yap* and *Taz* also produces isolated EA

Given that *Yap* deletion results in isolated EA and both Yap and Taz are key transcriptional effectors of the Hippo pathway, we investigated whether the simultaneous deletion of both *Yap* and *Taz* would worsen the foregut separation defect. We generated *Sox2-CreER;Yap^loxp/loxp^*;*Taz^loxp/loxp^* mutants and administered tamoxifen to pregnant females at E8.5. The mutant embryos were dead by E15.5 and exhibited facial abnormalities including a facial cleft and reduced eye size (Fig. 5A). At E12.5, similar to *Sox2-CreER;Yap^loxp/loxp^*mutants, deletion of *Yap* and *Taz* resulted in the formation of isolated EA (Fig. 5B). Together, these results further emphasize that Yap is required for tracheoesophageal separation, and that additional deletion of *Taz* did not appear to exacerbate the esophageal defect caused by *Yap* loss.

**Fig. 5.**
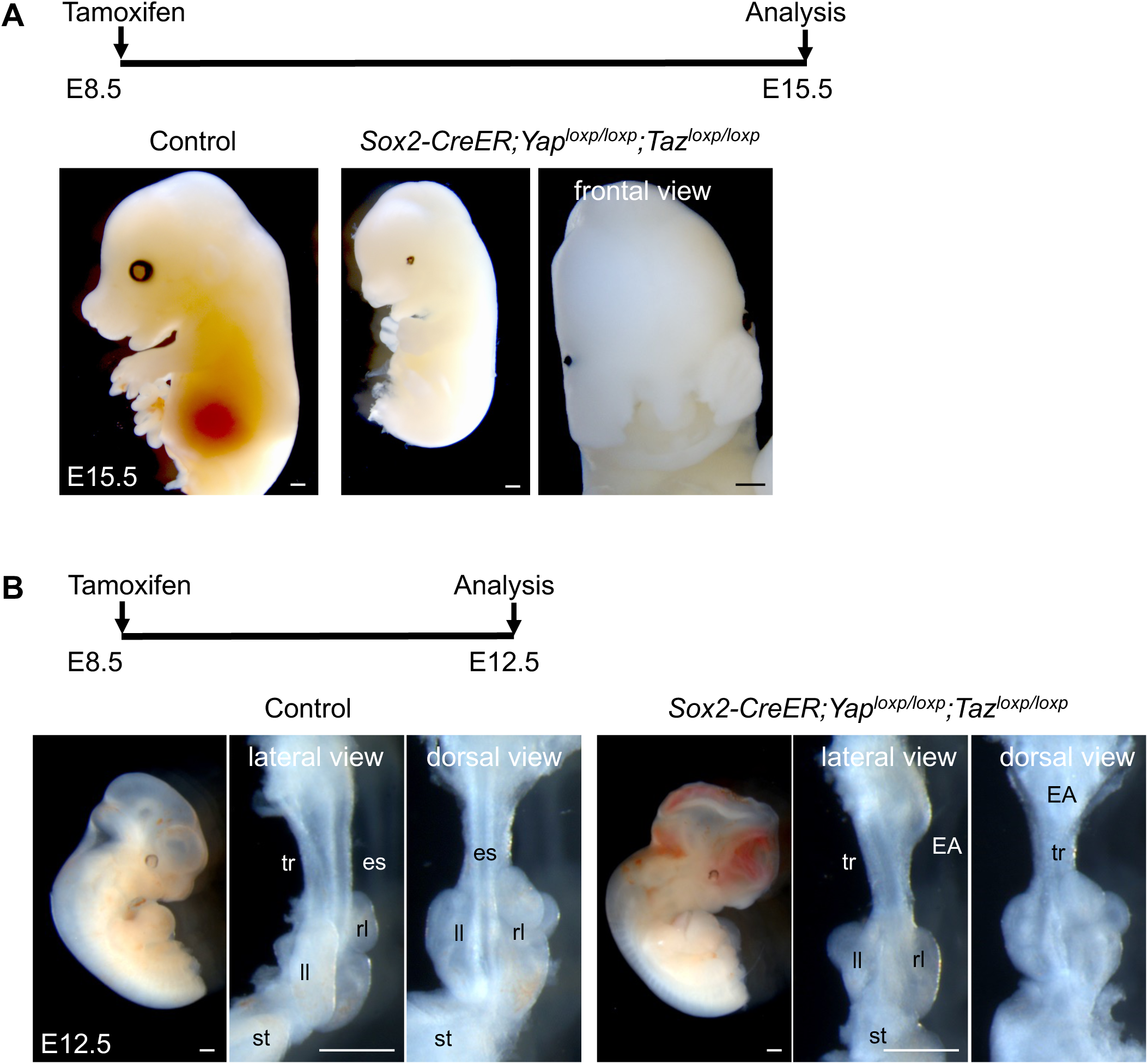
Combined deletion of *Yap* and *Taz* using *Sox2-CreER* produces isolated EA and severe craniofacial abnormalities. (A-B) Representative whole-mount images of control and *Sox2- CreER;Yap^loxp/loxp^;Taz^loxp/loxp^* mutant embryos (n=5). (A) *Yap* and *Taz* deletion mutant embryos die by E15.5 and show facial clefts and reduced eye size. (B) At E12.5, the mutants display isolated EA and cranial hemorrhage. EA, esophageal atresia; es, esophagus; ll, left lung; rl, right lung; st, stomach; tr, trachea. Scale bars: 1 mm.

In summary, our study identifies a critical role for Yap in tracheoesophageal separation. Our findings suggest that Yap regulates epithelial cells at the dorsoventral boundary of the anterior foregut, thereby contributing to the coordinated cellular processes underlying the separation of the developing respiratory and digestive tracts. Future studies should investigate the dynamic cellular behaviors at the epithelial intercalation site using live imaging and lineage-tracing approaches to characterize cell migration, intercalation, and rearrangement in real time. Furthermore, elucidating how Yap regulates these cellular processes and coordinates interactions between respiratory and esophageal epithelial cells will provide important mechanistic insights into dorsoventral boundary establishment and the pathogenesis of EA/TEF.

## Materials and Methods

### Mouse lines

*Sox2-CreER* (Arnold et al., 2011), *Yap^loxp/loxp^* (Schlegelmilch et al., 2011), and *Taz^loxp/loxp^*(Wang et al., 2016) mouse lines have been described previously. Mouse lines were maintained on a mixed C57BL/6 and 129SvEv background. The day of vaginal plug detection was designated as E0.5. Tamoxifen (Sigma, T5648) was prepared in sunflower seed oil (Sigma, S5007) at a stock concentration of 20 mg/ml. Pregnant females received a single intraperitoneal injection of tamoxifen (0.1 mg/g body weight) at E8.5. All animal studies were approved by the Columbia University Institutional Animal Care and Use Committee (IACUC).

### Tissue preparation and histology

Embryos or tissues were harvested in cold phosphate-buffered saline (PBS) and fixed overnight at 4°C in 4% paraformaldehyde in PBS. Following fixation, samples for paraffin sectioning were dehydrated through a graded series of increasing ethanol concentrations and then cleared with Histo-Clear. After embedding the specimens in paraffin, 7 µm sections were obtained from the blocks. For cryosectioning, the tissues were incubated in 15% sucrose in PBS until they sank and then transferred to 30% sucrose in PBS overnight. The samples were subsequently embedded in optimal cutting temperature (OCT) compound, and 8 µm sections were cut on a cryostat.

For Hematoxylin & Eosin staining, paraffin sections were first deparaffinized in Histo- Clear and then rehydrated through a graded series of ethanol. The sections were stained with Mayer’s Hematoxylin for 5 minutes, followed by a 10-minute wash with tap water. After counterstaining with Eosin for 3 minutes, the sections were dehydrated in ethanol solutions, cleared in Histo-Clear, and mounted with Permount Mounting Medium (Fisher Scientific).

### Immunofluorescence staining

For immunofluorescence staining, paraffin slides were deparaffinized, rehydrated, and then underwent heat-induced antigen retrieval with a citrate-based buffer (Vector Laboratories, H-3300). Cryosections were air-dried and washed in PBS. Slides were blocked for 1 hour at room temperature (RT) in 5% normal donkey serum solution in PBS with 0.3% Triton X-100. Slides were incubated with primary antibodies at 4°C overnight, and the following day, secondary antibodies were applied at a 1:500 dilution for 2 hours at RT. Primary antibodies were used as follows: acetylated α-tubulin (Sigma, T7451, 1:100), α-SMA (Sigma, A2547, 1:300), E-cadherin (Sigma, U3254, 1:500), Foxj1 (eBioscience, 14-9965-82, 1:200), Ki67 (Cell Signaling, 9129, 1:400), Krt8 (Developmental Studies Hybridoma Bank, TROMA-I, 1:100), Krt13 (Abcam, ab92551, 1:400), Nkx2.1 (Abcam, ab76013, 1:500), p63 (Cell Signaling, 13109, 1:400), Scgb1a1 (Santa Cruz Biotechnology, sc-365992, 1:200), Sftpc (Abcam, ab211326, 1:200), Sox2 (eBioscience, 14-9811-82, 1:400), Sox9 (R&D Systems, AF3075, 1:300), T1α (Developmental Studies Hybridoma Bank, 8.1.1, 1:200), Yap (Cell Signaling, 14074, 1:100). Alexa Fluor 488, 555, and 647-conjugated antibodies (Thermo Fisher Scientific, 1:500) were used as secondary antibodies. Nuclei were counterstained with DAPI (Sigma-Aldrich, D9542, 1:1,000).

Immunofluorescence images were acquired using a Zeiss LSM700 confocal laser scanning microscope (Carl Zeiss). Brightfield images were obtained with a Nikon SMZ1500 stereomicroscope (Nikon). H&E staining images were captured using a Leica DMi8 widefield microscope (Leica).

---

**Fig. S1.**
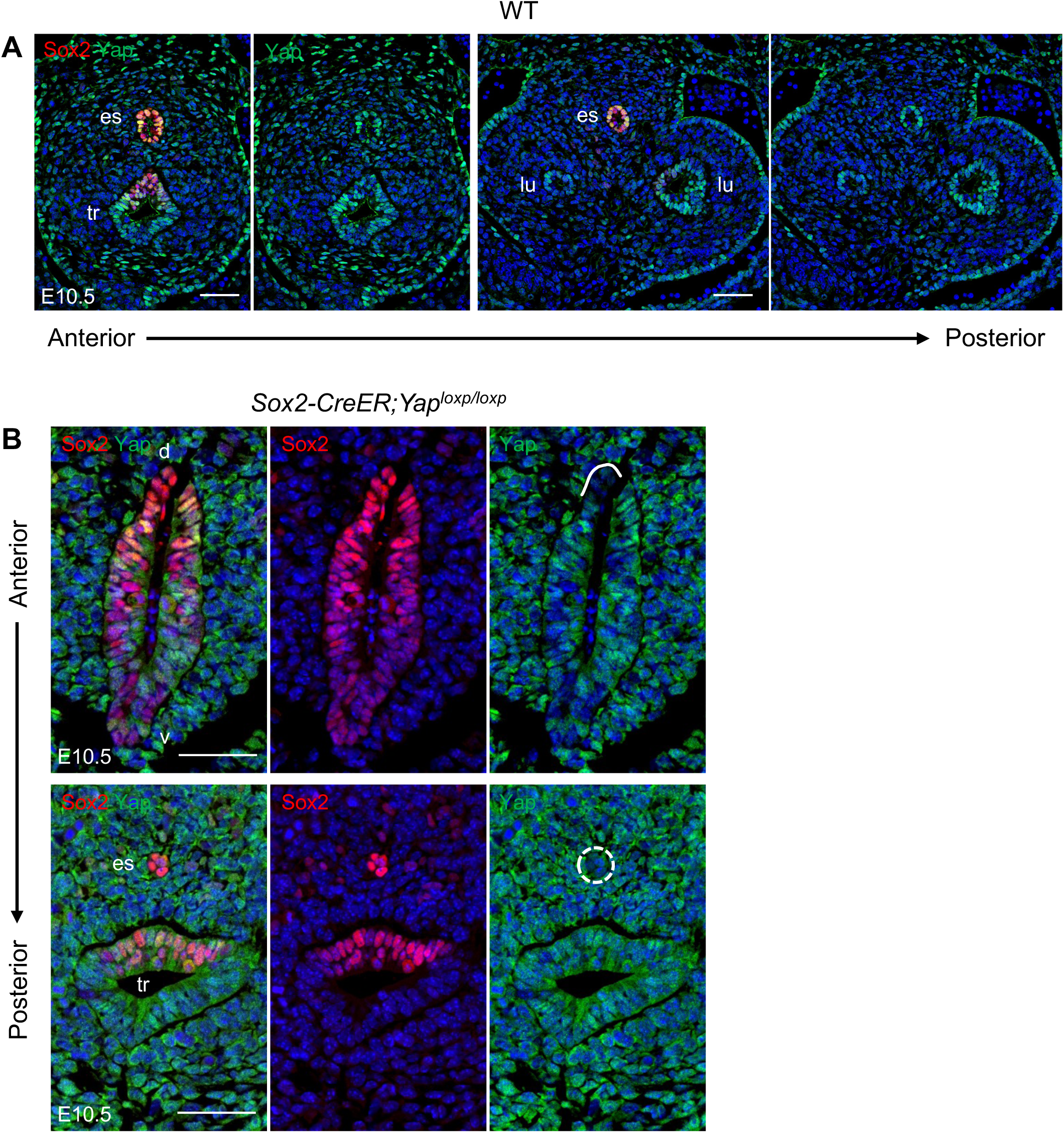
Yap expression is lost in the esophageal epithelium of *Yap* mutant foregut. (A) Yap is expressed in the epithelium of the esophagus, trachea, and lung buds in E10.5 wild-type embryos. (B) The absence of Yap expression in the dorsal foregut (solid line) and esophagus (dashed circle) of the *Sox2-CreER;Yap^loxp/loxp^* mutant at E10.5. d, dorsal; es, esophagus; lu, lung; tr, trachea; v, ventral; WT, wild type. Scale bars: 50 μm.

**Fig. S2.**
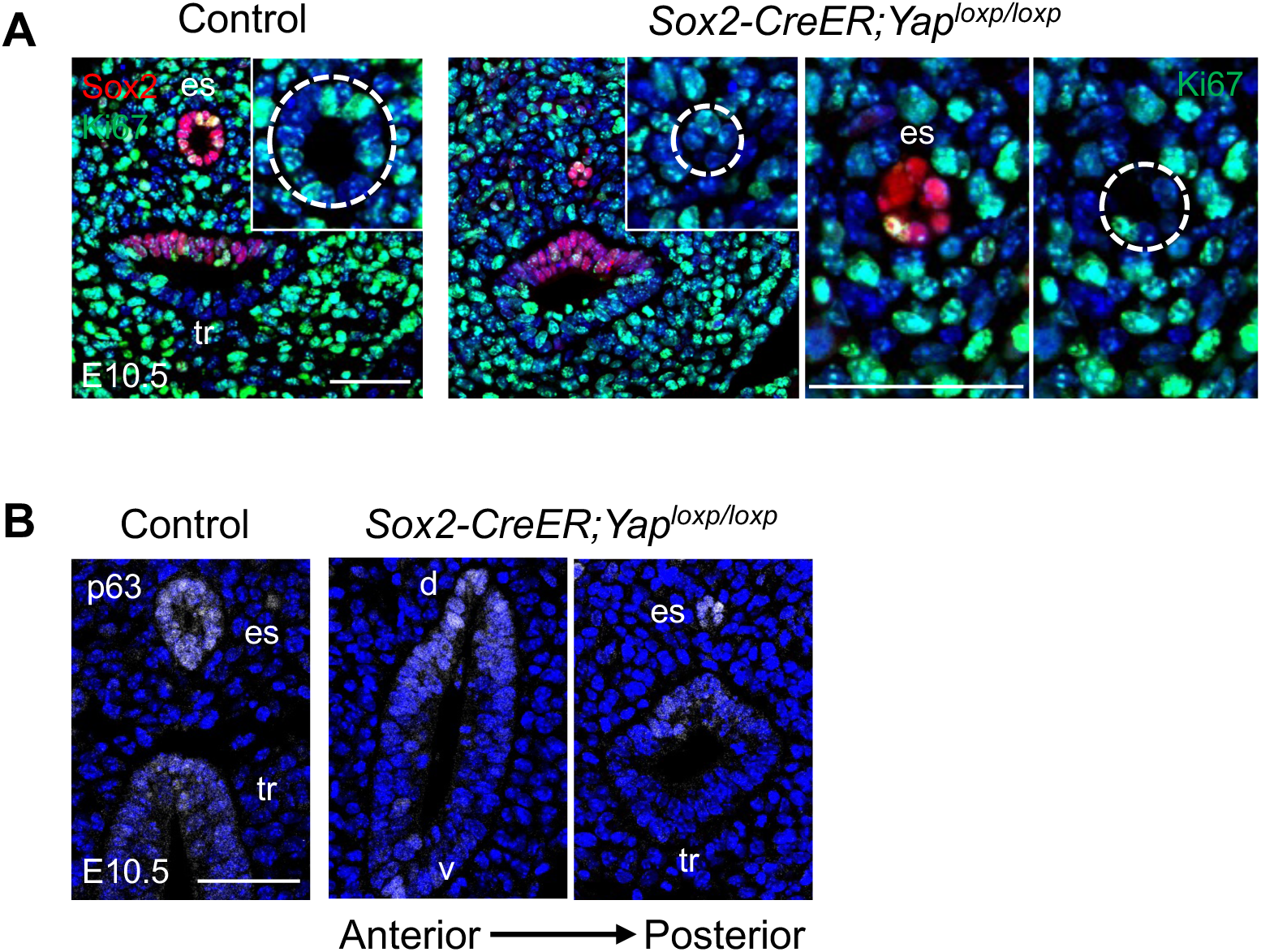
Esophageal progenitor cells are present in the *Yap* mutant foregut at E10.5. (A) Ki67^+^ cells are detected within the residual Sox2^+^ epithelial cells, although they are sparse in number (dashed circle). (B) p63 staining marks esophageal progenitor cells in the dorsal esophageal epithelium. d, dorsal; es, esophagus; tr, trachea; v, ventral. Scale bars: 50 μm.

**Fig. S3.**
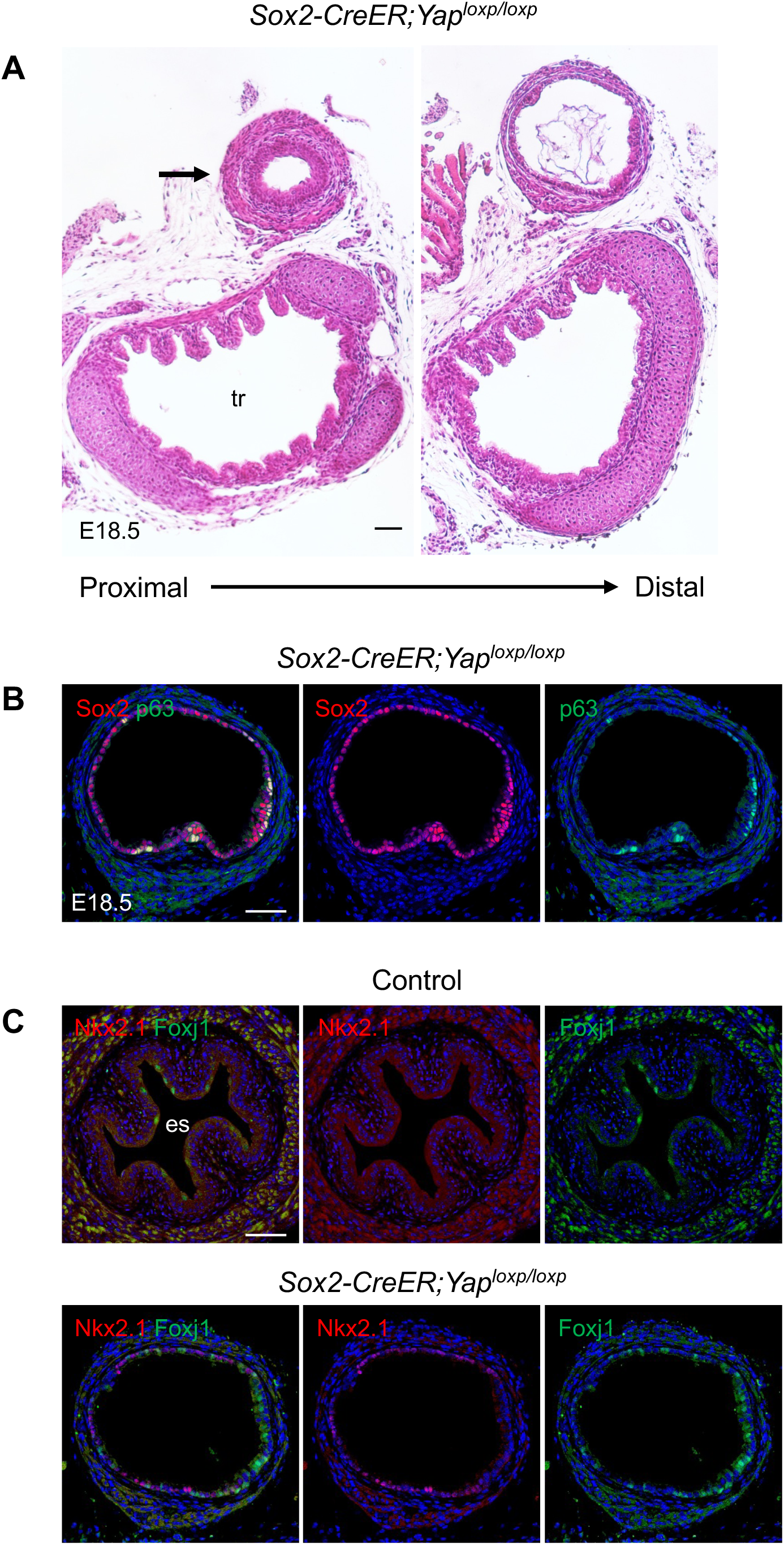
An ectopic epithelial tube expressing respiratory markers is present dorsal to the trachea in E18.5 *Sox2-CreER;Yap^loxp/loxp^*mutants. (A) An ectopic tube (arrow) adjacent to the trachea, composed of epithelial and mesenchymal layers but lacking cartilage upon *Yap* deletion. (B) Sox2 is expressed in most epithelial cells of the tube, whereas p63 is detected in a subset. (C) Some epithelial cells lining the tube express Nkx2.1 and Foxj1. Note that Nkx2.1 is not detected in the control esophageal epithelium. es, esophagus; tr, trachea. Scale bars: 50 μm.

## References

Arnold, K., Sarkar, A., Yram, M. A., Polo, J. M., Bronson, R., Sengupta, S., Seandel, M., Geijsen, N. and Hochedlinger, K. (2011). Sox2(+) adult stem and progenitor cells are important for tissue regeneration and survival of mice. Cell Stem Cell 9, 317–329.

Bailey, D. D., Zhang, Y., van Soldt, B. J., Jiang, M., Suresh, S., Nakagawa, H., Rustgi, A. K., Aceves, S. S., Cardoso, W. V. and Que, J. (2019). Use of hPSC-derived 3D organoids and mouse genetics to define the roles of YAP in the development of the esophagus. Development 146, dev178855.

Cardoso, W. V. and Lü, J. (2006). Regulation of early lung morphogenesis: questions, facts and controversies. Development 133, 1611–1624.

Domyan, E. T., Ferretti, E., Throckmorton, K., Mishina, Y., Nicolis, S. K. and Sun, X. (2011). Signaling through BMP receptors promotes respiratory identity in the foregut via repression of Sox2. Development 138, 971–981.

Geneviève, D., de Pontual, L., Amiel, J., Sarnacki, S. and Lyonnet, S. (2007). An overview of isolated and syndromic oesophageal atresia. Clin Genet 71, 392–399.

Goss, A. M., Tian, Y., Tsukiyama, T., Cohen, E. D., Zhou, D., Lu, M. M., Yamaguchi, T. P. and Morrisey, E. E. (2009). Wnt2/2b and beta-catenin signaling are necessary and sufficient to specify lung progenitors in the foregut. Dev Cell 17, 290–298.

Haight, C. (1944). Congenital Atresia of the Esophagus With Tracheoesophageal Fistula : Reconstruction of Esophageal Continuity by Primary Anastomosis. Ann Surg 120, 623–655.

Harris-Johnson, K. S., Domyan, E. T., Vezina, C. M. and Sun, X. (2009). beta-Catenin promotes respiratory progenitor identity in mouse foregut. Proc Natl Acad Sci U S A 106, 16287–16292.

Jacobs, I. J., Ku, W. Y. and Que, J. (2012). Genetic and cellular mechanisms regulating anterior foregut and esophageal development. Dev Biol 369, 54–64.

Kim, E., Jiang, M., Huang, H., Zhang, Y., Tjota, N., Gao, X., Robert, J., Gilmore, N., Gan, L. and Que, J. (2019). Isl1 Regulation of Nkx2.1 in the Early Foregut Epithelium Is Required for Trachea-Esophageal Separation and Lung Lobation. Dev Cell 51, 675–683.e4.

Kluth, D. and Fiegel, H. (2003). The embryology of the foregut. Semin Pediatr Surg 12, 3–9.

Krishnan, U. (2019). Eosinophilic esophagitis in esophageal atresia. Front. Pediatr. 7, 497.

Lei, Q. Y., Zhang, H., Zhao, B., Zha, Z. Y., Bai, F., Pei, X. H., Zhao, S., Xiong, Y. and Guan, K. L. (2008). TAZ promotes cell proliferation and epithelial-mesenchymal transition and is inhibited by the hippo pathway. Mol Cell Biol 28, 2426–2436.

Li, Y., Litingtung, Y., Ten Dijke, P. and Chiang, C. (2007). Aberrant Bmp signaling and notochord delamination in the pathogenesis of esophageal atresia. Dev Dyn 236, 746–754.

Litingtung, Y., Lei, L., Westphal, H. and Chiang, C. (1998). Sonic hedgehog is essential to foregut development. Nat Genet 20, 58–61.

Mahlapuu, M., Enerbäck, S. and Carlsson, P. (2001). Haploinsufficiency of the forkhead gene Foxf1, a target for sonic hedgehog signaling, causes lung and foregut malformations. Development 128, 2397–2406.

Malmström, K., Lohi, J., Lindahl, H., Pelkonen, A., Kajosaari, M., Sarna, S., Malmberg, L. P. and Mäkelä, M. J. (2008). Longitudinal follow-up of bronchial inflammation, respiratory symptoms, and pulmonary function in adolescents after repair of esophageal atresia with tracheoesophageal fistula. J Pediatr 153, 396–401.e1.

Minoo, P., Su, G., Drum, H., Bringas, P. and Kimura, S. (1999). Defects in tracheoesophageal and lung morphogenesis in Nkx2.1(-/-) mouse embryos. Dev Biol 209, 60–71.

Morin-Kensicki, E. M., Boone, B. N., Howell, M., Stonebraker, J. R., Teed, J., Alb, J. G., Magnuson, T. R., O’Neal, W. and Milgram, S. L. (2006). Defects in yolk sac vasculogenesis, chorioallantoic fusion, and embryonic axis elongation in mice with targeted disruption of Yap65. Mol Cell Biol 26, 77–87.

Motoyama, J., Liu, J., Mo, R., Ding, Q., Post, M. and Hui, C. C. (1998). Essential function of Gli2 and Gli3 in the formation of lung, trachea and oesophagus. Nat Genet 20, 54–57.

O’Rahilly, R. and Müller, F. (1984). Chevalier Jackson lecture. Respiratory and alimentary relations in staged human embryos. New embryological data and congenital anomalies. Ann Otol Rhinol Laryngol 93, 421–429.

Pan, D. (2010). The hippo signaling pathway in development and cancer. Dev Cell 19, 491–505.

Pepicelli, C. V., Lewis, P. M. and McMahon, A. P. (1998). Sonic hedgehog regulates branching morphogenesis in the mammalian lung. Curr Biol 8, 1083–1086.

Que, J., Choi, M., Ziel, J. W., Klingensmith, J. and Hogan, B. L. (2006). Morphogenesis of the trachea and esophagus: current players and new roles for noggin and Bmps. Differentiation 74, 422–437.

Que, J., Okubo, T., Goldenring, J. R., Nam, K. T., Kurotani, R., Morrisey, E. E., Taranova, O., Pevny, L. H. and Hogan, B. L. (2007). Multiple dose-dependent roles for Sox2 in the patterning and differentiation of anterior foregut endoderm. Development 134, 2521–2531.

Rodriguez, P., Da Silva, S., Oxburgh, L., Wang, F., Hogan, B. L. M. and Que, J. (2010). BMP signaling in the development of the mouse esophagus and forestomach. Development 137, 4171–4176.

Schlegelmilch, K., Mohseni, M., Kirak, O., Pruszak, J., Rodriguez, J. R., Zhou, D., Kreger, B. T., Vasioukhin, V., Avruch, J., Brummelkamp, T. R. and Camargo, F. D. (2011). Yap1 acts downstream of α-catenin to control epidermal proliferation. Cell 144, 782–795.

Spitz, L. (2007). Oesophageal atresia. Orphanet J Rare Dis 2, 24.

Torfs, C. P., Curry, C. J. R. and Bateson, T. F. (1995). Population-based study of tracheoesophageal fistula and esophageal atresia. Teratology 52, 220–232.

Wang, J., Xiao, Y., Hsu, C. W., Martinez-Traverso, I. M., Zhang, M., Bai, Y., Ishii, M., Maxson, R. E., Olson, E. N., Dickinson, M. E., Wythe, J. D. and Martin, J. F. (2016). Yap and Taz play a crucial role in neural crest-derived craniofacial development. Development 143, 504–515.

Warburton, D., Schwarz, M., Tefft, D., Flores-Delgado, G., Anderson, K. D. and Cardoso, W. V. (2000). The molecular basis of lung morphogenesis. Mech Dev 92, 55–81.

Wood, H. B. and Episkopou, V. (1999). Comparative expression of the mouse Sox1, Sox2 and Sox3 genes from pre-gastrulation to early somite stages. Mech Dev 86, 197–201.

Yan, R., Hoffmann, L. A., Oikonomou, P., Li, D., Lee, C., Gill, H. K., Mongera, A., Nerurkar, N. L., Mahadevan, L. and Tabin, C. J. (2025). Convergent flow-mediated mesenchymal force drives embryonic foregut constriction and splitting. Nat Commun 16, 10643.

Zhao, B., Wei, X., Li, W., Udan, R. S., Yang, Q., Kim, J., Xie, J., Ikenoue, T., Yu, J., Li, L., Zheng, P., Ye, K., Chinnaiyan, A., Halder, G., Lai, Z. C. and Guan, K. L. (2007). Inactivation of YAP oncoprotein by the Hippo pathway is involved in cell contact inhibition and tissue growth control. Genes Dev 21, 2747–2761.

Zhao, B., Ye, X., Yu, J., Li, L., Li, W., Li, S., Yu, J., Lin, J. D., Wang, C. Y., Chinnaiyan, A. M., Lai, Z. C. and Guan, K. L. (2008). TEAD mediates YAP-dependent gene induction and growth control. Genes Dev 22, 1962–1971.

Zhao, B., Li, L., Tumaneng, K., Wang, C. Y. and Guan, K. L. (2010). A coordinated phosphorylation by Lats and CK1 regulates YAP stability through SCF(beta-TRCP). Genes Dev 24, 72–85.

